# IFT20 is critical for early chondrogenesis during endochondral ossification

**DOI:** 10.1101/2021.06.25.449945

**Authors:** Hiroyuki Yamaguchi, Megumi Kitami, Karin H. Uchima Koecklin, Li He, Jianbo Wang, Daniel S. Perrien, William R. Lagor, Yoshihiro Komatsu

**Affiliations:** Department of Pediatrics, McGovern Medical School, UTHealth, Houston, TX 77030; Division of Endocrinology, Metabolism, and Lipids, Department of Medicine, Emory University, Atlanta, GA 30232; Department of Molecular Physiology and Biophysics, Baylor College of Medicine, Houston TX 77030; Graduate Program in Genetics & Epigenetics, The University of Texas MD Anderson Cancer Center UTHealth Graduate School of Biomedical Sciences, Houston, TX 77030

**Keywords:** Cartilage, Cilia, FGF, Intraflagellar transport, Mice

## Abstract

Ciliogenic components, such as the family of intraflagellar transport (IFT) proteins, are recognized to play key roles in endochondral ossification, a critical process to form most bones. However, it remains unclear how each IFT protein performs its unique function to regulate endochondral ossification. Here, we show that intraflagellar transport 20 (IFT20) is required for early chondrogenesis. Utilizing three osteochondrocyte lineage-specific Cre mice (*Prx1-Cre*, *Col2-Cre* and *Aggrecan-CreER^T2^*), we deleted *Ift20* to examine its function. While chondrocyte-specific *Ift20* deletion with *Col2-Cre* or *Aggrecan-CreER^T2^* drivers did not cause overt skeletal defects, mesoderm-specific *Ift20* deletion using *Prx1-Cre* (*Ift20:Prx1-Cre*) resulted in shortened limb outgrowth. Although primary cilia were not formed in *Ift20:Prx1-Cre* mice, ciliary Hedgehog signaling was only moderately affected. Interestingly, loss of *Ift20* lead to upregulation of *Fgf18* expression resulting in ERK1/2 activation and sustained *Sox9* expression, thus preventing endochondral ossification. Inhibition of enhanced phospho-ERK1/2 activation partially rescued defective chondrogenesis in *Ift20* mutant cells, supporting an important role for FGF signaling. Our findings demonstrate a novel mechanism of IFT20 in early chondrogenesis during endochondral ossification.

## Introduction

Endochondral ossification is an essential process which forms the majority of bones^1,2^. Unlike intramembranous ossification, which consists of direct conversion of mesenchymal cells to osteoblasts, in endochondral ossification mesenchymal cells differentiate into chondrocytes that create a cartilage template. Then later in development osteoblasts replace this cartilage with bone^3,4^. In past decades, tremendous progress has been made toward understanding the molecular mechanisms of endochondral ossification^5,6^. These studies have identified genetic and growth factor signaling pathways that are tightly linked to govern spatiotemporal development of endochondral ossification. For example, transforming growth factor-β/bone morphogenetic protein (TGF-β/BMP), Wnt and Hedgehog signaling interact with each other to control proliferation and survival of oesteo-chondrogenic progenitor cells^7–9^. However, it remains elusive how growth factor signals are precisely transduced into these cells to control endochondral ossification.

Primary cilia are microtubule-based antenna-like organelles that project from the surface of vertebrate cells^10–12^. Previously we reported, along with others, that primary cilia play a pivotal role in regulating developmental signals, such as Hedgehog, TGF-beta, and platelet-derived growth factor (PDGF)^11,13^. Because primary cilia participate in controlling multiple developmental pathways, a better understanding of the regulation of ciliogenesis has become increasingly important as ciliary defects continue to be linked to numerous diseases, collectively called “ciliopathies”^14^. These findings suggest that dysfunction and/or altered formation of cilia may directly contribute to the pathogenesis of skeletal diseases.

A fundamental role of intraflagellar transport (IFT) complex is to assemble cilia^10,15^. In humans, mutations in ciliary genes frequently result in skeletal abnormalities^16–19^. There are several skeletal ciliopathies identified such as Verma-Naumoff syndrome, Majewski syndrome, Jeune syndrome, and Ellis-van Creveld syndrome^17,20^. Similar to these conditions, loss of IFT component function also results in abnormalities in skeletogenic cell proliferation, survival and differentiation in mice^21–24^. For example, IFT88, a component of the IFT-B complx, in the mesenchyme is important for embryonic limb outgrowth^23^. KIF3A, a subunit of the Kinesin II motor complex required for IFT, is critical for postnatal growth plate development, and formation of cranial base and synchondroses^22,25^. IFT80 is indispensable for cartilage and bone development in both embryonic and postnatal stages^26^. However, the roles played by other IFTs in bone formation and regulating the process of endochondral ossification remains unclear. Additionally, while ciliary Hedgehog signaling has been found to be an important process involved in skeletal ciliopathies, the phenotypes observed in IFT mutants cannot be solely explained by the alteration of ciliary Hedgehog signaling^27,28^. Therefore, it is important to explore whether other signaling pathway(s) are involved in the etiology of skeletal ciliopathies in IFT mutants.

In this study, we examine the function of IFT20, a component of the IFT-B complex, during endochondral ossification. Our findings demonstrate that IFT20 is required for early chondrogenesis, which has not been characterized before in endochondral ossification.

## Materials and methods

### Animals

*Ift20*-floxed mice^29^, *Prx1-Cre* mice^30^, and *Aggrecan-CreER^T2^* mice^31^ were obtained from the Jackson Laboratory. *Col2-Cre* mice^32^ were obtained from Dr. Florent Elefteriou (Baylor College of Medicine). These mice were maintained on a mixed genetic background. The Animal Welfare Committee, and the Institutional Animal Care and Use Committee of the McGovern Medical School approved the experimental protocol (AWC-18-0137).

### Histological analysis, skeletal staining and immunostaining

H&E, Safranin-O, Picrosirius red staining and von Kossa staining were carried out using standard methods^33^. Staining of long bones with Alizarin red and Alcian blue was carried out as previously described^33^. Immunofluorescent staining and TUNEL assays were performed as previously described^34^. Anti-acetylated tubulin (Sigma; T6793, 1:1,000), anti-gamma tubulin (Sigma; T5326, 1:1,000), anti-phospho Histone H3 (Millipore; #06-570, 1:100), anti-SOX9 (Santa cruz biotechnology; sc-20095, 1:100), anti-phospho ERK1/2 (Cell signaling technology; #4376, 1:100) antibodies were used for immunostaining. Slides were imaged under an Olympus FluoView FV1000 laser scanning confocal microscope using the software FV10-ASW Viewer (version 4.2).

### Section *in situ* hybridization

A digoxygenin-labelled RNA probe was generated by *in vitro* transcription (Sigma-Aldrich). *In situ* hybridization was performed according to standard procedures^35^.

### Micromass culture

Micromass culture was carried out using a standard method^36^. Briefly, mesenchymal cells were harvested from *Ift20^flox/flox^* limb buds at E11.5. After 0.25% trypsin treatment at 37°C for 10min, 15μl spots of cell suspension (2.5×10^5^ cells/spot) were plated onto 4-well culture dishes (Thermo Fisher Scientific; 176740) and cultured using Advanced DMEM medium (Sigma-Aldrich; 12491-023) containing 10% fetal bovine serum (Sigma-Aldrich; F4135), 2% L-glutamine (Sigma-Aldrich; G7513), and 1% Penicillin-Streptomycin (Sigma-Aldrich; P4333). To disrupt *Ift20*, adenovirus-Cre (5×10^9^ pfu, Vector lab, Baylor College of Medicine) was transduced in *Ift20^flox/flox^* limb mesenchymal cells at 37°C for overnight. As a control, adenovirus-empty vector was used. To inhibit the phospho-ERK1/2 activity, U0126 (Cell Signaling technology; 9903) was used. DMSO-vehicle solution was used as a control.

### Quantitative real-time RT-PCR

Total RNA was extracted using TRIzol Reagent (Thermo Fisher Scientific; 15596-026). Quantitative RT-PCR was carried out using the SsoAdvanced Universal SYBR Green Supermix (Bio-Rad; 1725274). The conditions for qRT-PCR were 95°C for 2 min, 95°C for 5 sec, and 60°C for 30 sec, for 40 cycles. The sequences of each primer are listed (Supplemental Table 1). Data were normalized to *Gapdh* levels and quantified using the 2^−ΔΔCT^ method.

### Western blotting analysis

Cell lysates were separated by SDS-PAGE (Bio-Rad; 4561036). Anti-IFT20 (Proteintech; 13615-AP, 1:1,000), anti-GAPDH (Cell Signaling technology; #2118, 1:5,000), anti-GLI1 (R&D systems; AF3455, 1:500), anti-SOX9 (Santa cruz biotechnology; sc-20095, 1:1,000), anti-phospho ERK1/2 (Cell Signaling technology; #4376, 1:1,000), anti-ERK1/2 (Cell Signaling technology; #4695, 1:1,000), anti-active β-Catenin (Cell signaling technology; #8814, 1:1,000), anti-β-Catenin (BD Biosciences; #610153, 1:1,000), anti-rabbit IgG HRP-conjugate (Cell Signaling technology; #7074, 1:5,000), anti-mouse IgG HRP-conjugate (Cell Signaling technology; #7076, 1:5,000) and anti-goat IgG HRP-conjugate (Invitrogen; #PA1-28664, 1:2,000) antibodies were used for western blotting. The Clarity Max ECL Substrate (Bio-Rad; 1705061) was used for chemiluminescent detection on a Chemidoc XRS+ imaging system (Bio-Rad), and the signals were quantified using the Image Lab Version 6.0 software (Bio-Rad) and the image-J software.

### Statistical analysis

A two-tailed Student’s *t* test was used for comparisons between two groups. A *p* value of less than 0.05 was considered statistically significant.

## Results and Discussion

### Mesoderm-specific disruption of *Ift20* results in a shorter limb development

To examine the function of IFT20 during endochondral ossification, *Ift20* floxed mice and *Prx1-Cre* mice were used to delete *Ift20* in a mesodermal cell-specific manner (“*Ift20:Prx1-Cre*” hereafter). Around the age of weaning, the locomotion of *Ift20:Prx1-Cre* mice is defective (Supplemental video 1). This is likely due to the formation of shorter limbs in *Ift20:Prx1-Cre* mice (Fig. 1A). Skeletal staining revealed short-limbed dwarfism with polydactyly in *Ift20:Prx1-Cre* mice (Fig. 1B). Polydactyly is a common abnormality frequently observed in ciliopathies and an expected result of altered ciliary Hedgehog signaling^23,37,38^. In this paper, we focus specifically on investigating the molecular basis for the defective limb phenotypes.

**Fig. 1.**
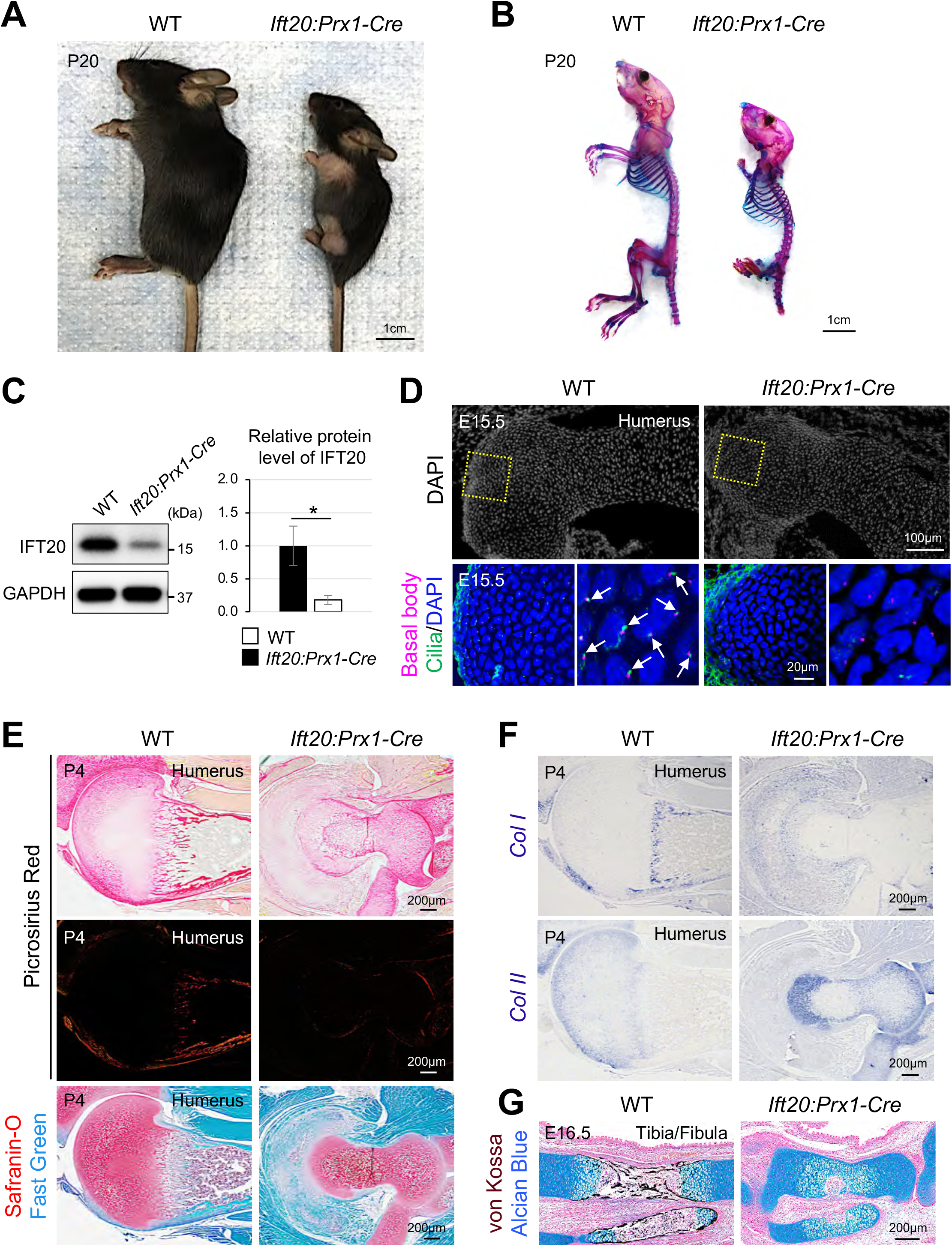
Mesoderm-specific disruption of *Ift20* results in a shorter limb development. **(A, B)** Whole mount view and skeletal staining analysis of wild-type (WT) and *Ift20:Prx1-Cre* mice at postnatal day (P) 20. **(C)** Western blotting analysis using tissue lysates from forelimbs at embryonic day (E) 15.5. **(D)** Immunohistochemical analysis to examine the presence of primary cilia. **(E)** Picrosirius Red staining and Safranin-O staining using humerus tissue section. **(F)** Gene expression analysis of *Col I* and *Col II* by section in situ hybridization. **(G)** von Kossa and Alcian blue double staining. Data in (C) are represented as mean ± SD, n = 3 in each group. *p<0.05.

Compared with wild-type controls (“WT” hereafter), the body size is significantly smaller in *Ift20:Prx1-Cre* mice (Fig. 1A, B). However, this growth retardation is possibly a secondary phenotype. Due to mobility issues, *Ift20:Prx1-Cre* mice have difficulties accessing the dam, and may not nurse effectively compared to WT littermates. Detailed morphological analysis revealed that these characteristic phenotypes were evident by embryonic day (E) 18.5 (Supplemental Fig. 1). Western blotting analysis using embryonic limb lysates confirmed that IFT20 is knocked-out in a subset of cells in *Ift20:Prx1-Cre* mice (Fig. 1C). Confocal microscopy analysis revealed that while the basal body was present, no primary cilia are formed in *Ift20:Prx1-Cre* humerus (Fig. 1D).

At P4, compared with WT, picrosirius red staining is weaker in *Ift20:Prx1-Cre* mice (Fig. 1E, upper and middle panels). This suggests that production of type I collagen was severely attenuated in *Ift20:Prx1-Cre* mice. To examine whether the reduction of type I collagen was due to altered endochondral ossification, cartilage formation was also examined. In *Ift20:Prx1-Cre* mice, the entire area of the humerus stains positively with Safranin-O (Fig. 1E, lower panel). This suggests that cartilage template was not appropriately replaced by osteoblasts in *Ift20:Prx1-Cre* mice. *In situ* hybridization analysis further confirmed that, compared with WT, type I collagen expression was decreased, but the expression of type II collagen was sustained in *Ift20:Prx1-Cre* mice at P4 (Fig. 1F). To determine whether the initial conversion process from cartilage template to osteoblasts was affected, embryonic hindlimb section was examined by double staining of von Kossa and Alcian blue. At E16.5, the process of bone mineralization was hampered in *Ift20:Prx1-Cre* mice (Fig. 1G). These results suggest that IFT20 plays a pivotal role during endochondral ossification.

### Chondrocyte-specific disruption of *Ift20* does not cause bone and cartilage defects

Endochondral ossification is a series of developmental processes initiated by appropriate chondrogenesis: (i) differentiation of mesenchymal cells to chondrogenic precursors, (ii) differentiation of chondrogenic precursors to chondrocytes, and (iii) maturation of chondrocytes. Depending on these developmental stages, each ciliary gene may function differently. To clarify the role of IFT20 at each chondrogenic stage, *Col2-Cre* mice was used to delete *Ift20* (“*Ift20:Col2-Cre*” hereafter). Unlike *Kif3a:Col2-Cre* mutant mice^22,25^, *Ift20:Col2-Cre* mice did not survive after birth. The exact cause of lethality is unknown. Chondrocyte-specific disruption of *Ift20* results in loss of primary cilia (Fig. 2A), suggesting that *Ift20* was efficiently deleted in *Ift20:Col2-Cre* chondrocytes. We noted that primary cilia in the perichondrial layer are intact in *Ift20:Col2-Cre* mice (Fig. 2A). This observation is consistent with previous reports that *Col2-Cre* does not effectively target the perichondrium^22,32^. Unlike *Ift80:Col2-Cre* mutant mice^26^, growth morphology of both forelimb and hindlimb was comparable between WT and *Ift20:Col2-Cre* mice (Fig. 2B). Histological analysis confirmed that there were no overt skeletal defects in *Ift20:Col2-Cre* mice (Fig. 2C).

**Fig. 2.**
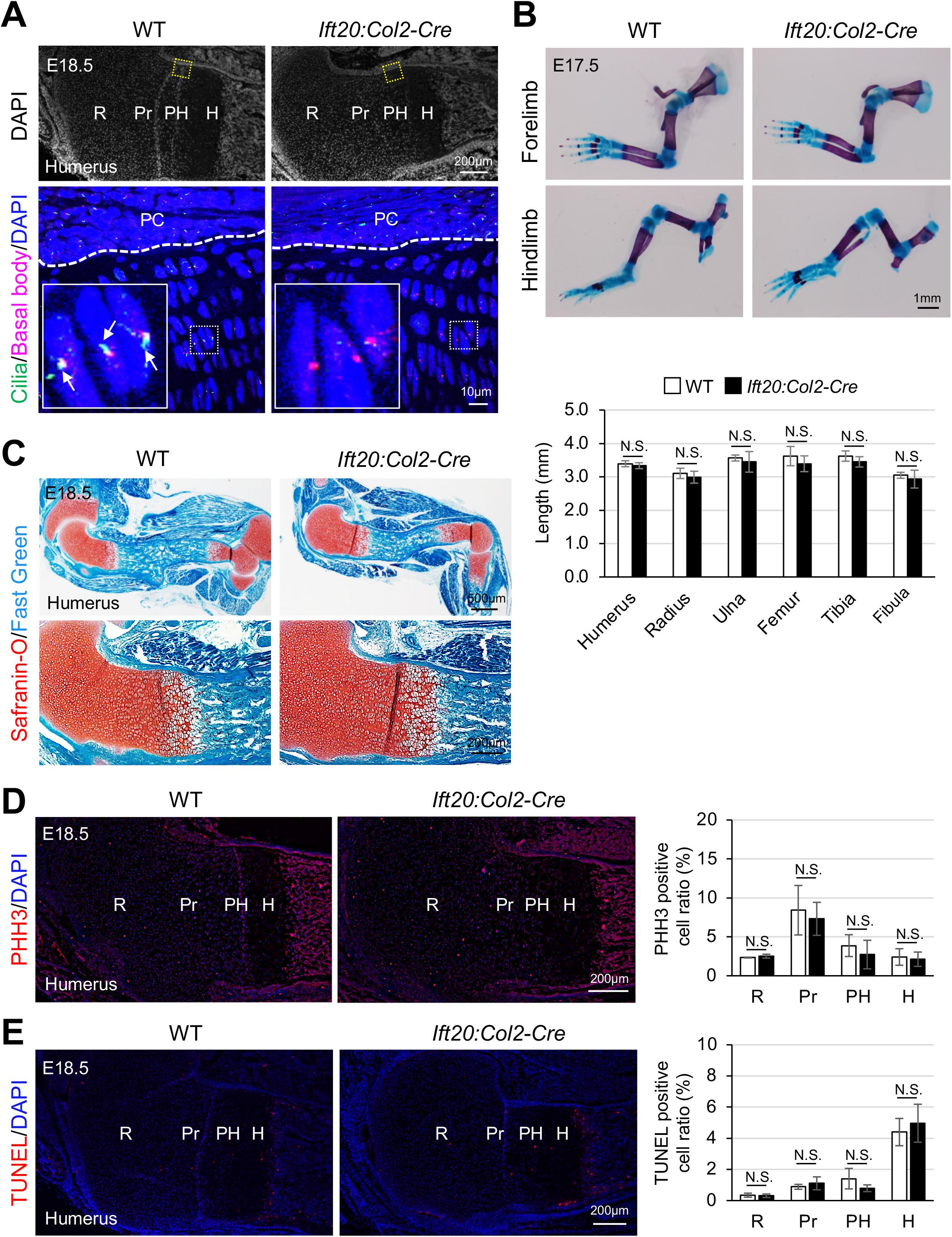
Chondrocyte-specific disruption of *Ift20* does not cause limb defects. **(A)** Immunohistochemical analysis to examine the presence of primary cilia. R, resting; Pr, proliferating; PH, pre-hypertrophic; H, hypertrophic; PC, perichondrium. Arrows indicate the primary cilia. **(B)** Skeletal staining analysis and quantification of the length of forelimbs and hindlimbs in WT and *Ift20:Col2-Cre* mice at E17.5. **(C)** Safranin-O staining. **(D)** Immunohistochemical analysis for pHH3 (magenta) and quantification analysis using humerus tissues. R, resting; Pr, proliferating; PH, pre-hypertrophic; H, hypertrophic. **(E)** TUNEL assay (magenta) and quantification analysis in humerus tissues. R, resting; Pr, proliferating; PH, pre-hypertrophic; H, hypertrophic. Data in (B), (D) and (E) are represented as mean ± SD, n = 3 in each group. *p<0.05, N.S., not significant.

To examine whether the loss of *Ift20* in *Col2-Cre* labeled chondrocytes affects cell viability, cell proliferation activity and survivability were examined by phospho-H3 (PHH3) immunostaining and TUNEL assay. As expected, chondrogenic proliferation and survival were comparable between WT and *Ift20:Col2-Cre* mice (Fig. 2D, E). To further examine whether IFT20 is predominantly involved in the regulation of early chondrogenesis, *Aggrecan-CreER^T2^* mice were used to examine the role of IFT20 during chondrocyte maturation (“*Ift20:Aggrecan-CreER^T2^*” hereafter). Similar to *Ift20:Col2-Cre* mice, no overt forelimb and/or hindlimb abnormalities were found in *Ift20:Aggrecan-CreER^T2^* mice (Supplemental Fig. 2). These results suggest that IFT20 is indispensable for early chondrogenesis but dispensable for the stages of chondrocyte differentiation and maturation during endochondral ossification.

### Ciliary Hedgehog signaling is moderately affected in *Ift20:Prx1-Cre* mice

Because *Ift20:Prx1-Cre* mice displayed shortened limb outgrowth, we hypothesize whether this defect is attributed to the decreased cell proliferation and/or increased cell death. To address these questions, cell proliferation and survivability were examined by immunohistochemistry and TUNEL assay. Compared with WT, *Ift20:Prx1-Cre* mice showed decreased cell proliferation activity while increased cell death (Fig. 3A, B), suggesting both chondrogenic cell proliferation and survival were affected.

**Fig. 3.**
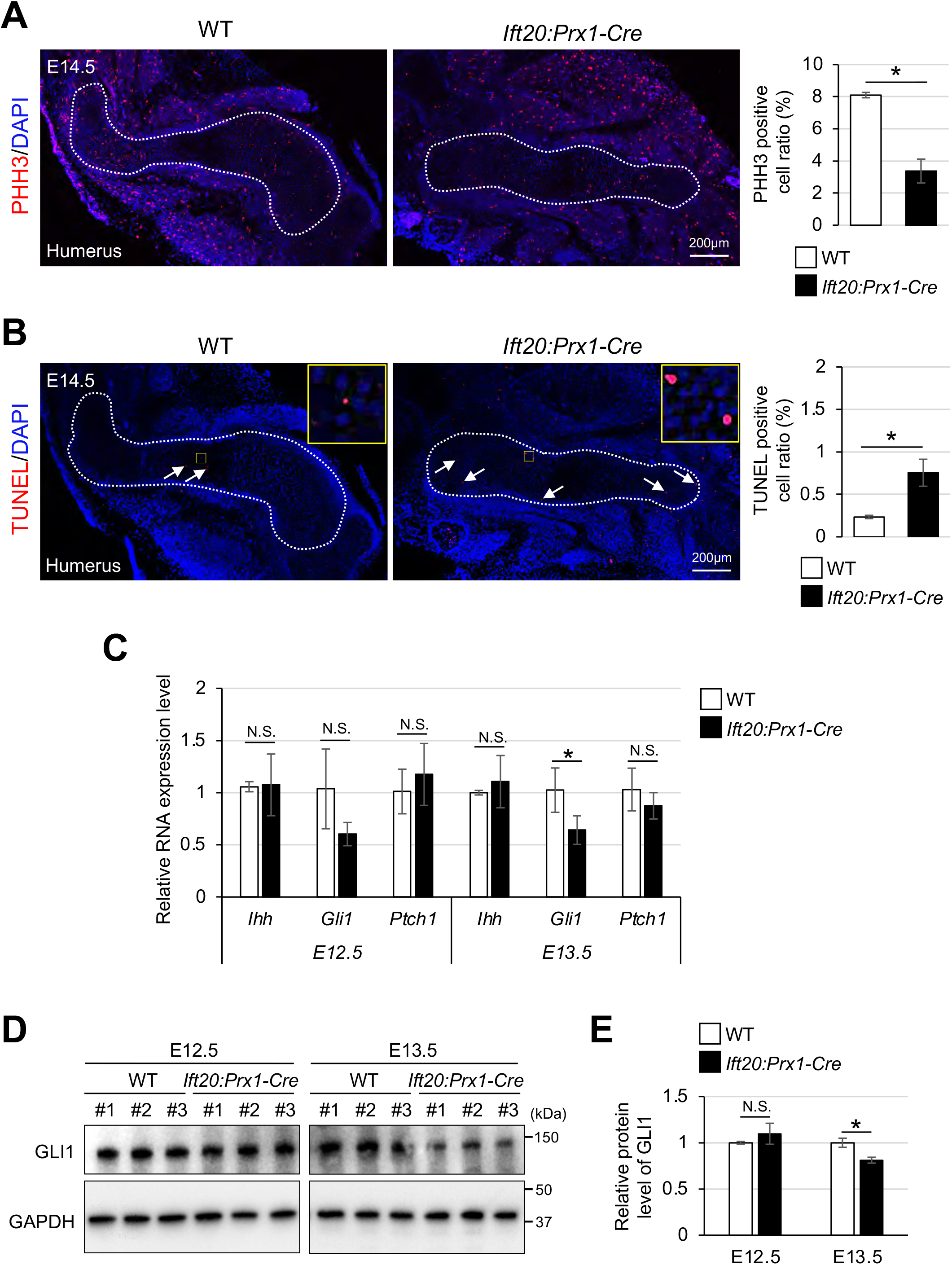
Ciliary Hedgehog signalling is moderately affected in *Ift20:Prx1-Cre* mice. **(A)** Immunohistochemical analysis for pHH3 (magenta) and quantification analysis using humerus tissues. **(B)** TUNEL assay (magenta) and quantification analysis in humerus tissues. Arrows show TUNEL positive signals. **(C)** qRT-PCR analysis for *Ihh*, *Gli1* and *Ptch1* using cell lysates from forelimb tissues. **(D)** Western blotting analysis for GLI1 using cell lysates from forelimb tissues. **(E)** Quantification analysis of GLI1 production from (E). Data in (A), (B), (C) and (E) are represented as mean ± SD, n = 3 in each group. *p<0.05, N.S., not significant.

Multiple studies have demonstrated that ciliary hedgehog signaling is required for proper bone growth^39–41^. For example, mesoderm-specific *Ift88* mutant mice and chondrocyte-specific *Ift80* mutant mice show the reduced Hedgehog signaling and leading to shorter limb development^23,26^. Therefore, it is reasonable to predict that the attenuation of ciliary Hedgehog signaling may cause abnormal limb outgrowth in *Ift20:Prx1-Cre* mice. To test this hypothesis, forelimb tissues (excluding limb buds) were isolated, then examined for gene expression related to Hedgehog signaling. Genes such as *Indian hedgehog* (*Ihh*), *Gli1* and *Ptch1* were quantified using qRT-PCR at E12.5-E13.5 when chondrogenic precursors start to differentiate. Among these Hedgehog signaling-associated genes, *Gli1* expression is slightly decreased in *Ift20:Prx1-Cre* mice (~E13.5) (Fig. 3C). GLI1 western blotting further confirmed that ciliary Hedgehog signaling is moderately affected in *Ift20:Prx1-Cre* mice (Fig. 3D, E). These observations are important because mesoderm-specific *Ift88* mutant mice develop similar limb outgrowth defects seen in *Ift20:Prx1-Cre* mice, but *Ift88* mutant’s abnormalities are attributed to the abrogated Hedgehog signaling (i.e., reduction of *Ihh* expression, no expression of *Ptch1* or *Gli1*)^23^. These results suggest that each ciliary component may have their own unique role to transduce signals during endochondral ossification. Therefore, in addition to the moderately altered Hedgehog signaling, other signaling cascade(s) may be involved in the pathology of *Ift20:Prx1-Cre* mice.

### Upregulation of FGF signaling prevents chondrogenesis in *Ift20:Prx1-Cre* mice

During skeletal development, fibroblast growth factor (FGF) signaling is another of the key regulators to coordinate endochondral ossification^2,4^. In the early embryonic stages of skeletal development, FGF signaling has promitogenic activity in chondrocytes. On the other hand, during late stages of skeletal growth, FGF signaling functions to inhibit chondrogenesis, preventing their differentiation to prehypertrophic and hypertrophic chondrocytes^42^. This paradoxical FGF signaling activity explains how the alteration of FGF signaling causes defective skeletal growth, frequently observed in skeletal dwarfism^43,44^. Because *Ift20:Prx1-Cre* mice showed only modest alterations of ciliary Hedgehog signaling (Fig. 3C-E), we hypothesized that aberrant FGF signaling may explain the defective limb outgrowth. To test this hypothesis, the levels of phospho-ERK1/2 (p-ERK1/2), a downstream target of FGF signaling, were examined by immunohistochemistry. In WT humerus, p-ERK1/2 signaling was mainly detected in the perichondrium, while p-ERK1/2 was highly activated in all cartilaginous cells in *Ift20:Prx1-Cre* mice (Fig. 4A). This p-ERK1/2 activation was also observed in *Ift20:Prx1-Cre* mice at P4 (Supplemental Fig. 3), suggesting that aberrant p-ERK1/2 activation causes abnormal chondrogenesis via suppressing hypertrophic differentiation. To examine whether the enhanced p-ERK1/2 activation is due to the upregulation of FGF ligands and/or its receptors in *Ift20:Prx1-Cre* mice, we examined expression of these genes by qRT-PCR analysis. Among FGF signaling-related genes, compared with WT, *Fgf18* expression was upregulated in *Ift20:Prx1-Cre* mice (Fig. 4B). To examine how the aberrant FGF signaling results in altered chondrogenesis, *Sox9* expression, a key regulator of chondrogenesis, was examined. During the early chondrogenesis stage (~E12.5), the levels of *Sox9* expression was comparable between WT and *Ift20:Prx1-Cre* mice (Fig. 4C). However, *Sox9* expression was increased during chondrogenesis differentiation (~E13.5) in *Ift20:Prx1-Cre* mice (Fig. 4C, D). *Sox9* expression was still sustained when osteogenesis initiates in *Ift20:Prx1-Cre* mice (~E15.5) (Fig. 4E). These observations are consistent with the evidence that overexpression of *Sox9* leads to delayed hypertrophy during chondrocyte maturation^45,46^. To examine whether augmentation of FGF pathway is involved in defective chondrogenesis in *Ift20:Prx1-Cre* mice, chondrocyte micromass culture was performed. After harvesting limb mesenchymal cells from *Ift20^flox/flox^* mice at E11.5, *Ift20* was deleted by infecting cells with an adenovirus that constitutively expresses Cre. Consistent with *in vivo* observation (Fig. 4A), p-ERK1/2 was highly activated in *Ift20* mutant cells (Fig. 4F). To suppress levels of p-ERK1/2 the small molecule inhibitor of MEK1/2 (U0126) was used. Treatment of U0126 suppressed the aberrant p-ERK1/2 activation in *Ift20* mutant cells (Fig. 4F), and inhibition of p-ERK1/2 partially rescued the defective cartilaginous nodule formation in *Ift20* mutant cells (Fig. 4G). These results suggest that while we found a moderate alterations in Hedgehog signaling, there was a much greater change in FGF-ERK1/2 signaling in *Ift20* mutants. Our data suggests that disruption of IFT20 increases the expression of *Fgf18* that leads to p-ERK1/2 activation, thus sustaining SOX9 expression and suppressing hypertrophic differentiation in *Ift20:Prx1-Cre* mice (Fig. 4H). Because FGF signaling can activate Wnt/β-catenin signaling and suppresses hypertrophic differentiation^47,48^, we also hypothesized that FGF-Wnt/β-catenin signaling pathway may cause shorter limb formation in *Ift20:Prx1-Cre* mice. However, the levels of both active β-catenin and total β-catenin were comparable between WT and *Ift20:Prx1-Cre* mice (Supplemental Fig. 4). This suggests that activation of the FGF signaling pathway may predominantly hamper endochondral ossification process in *Ift20:Prx1-Cre* mice.

**Fig. 4.**
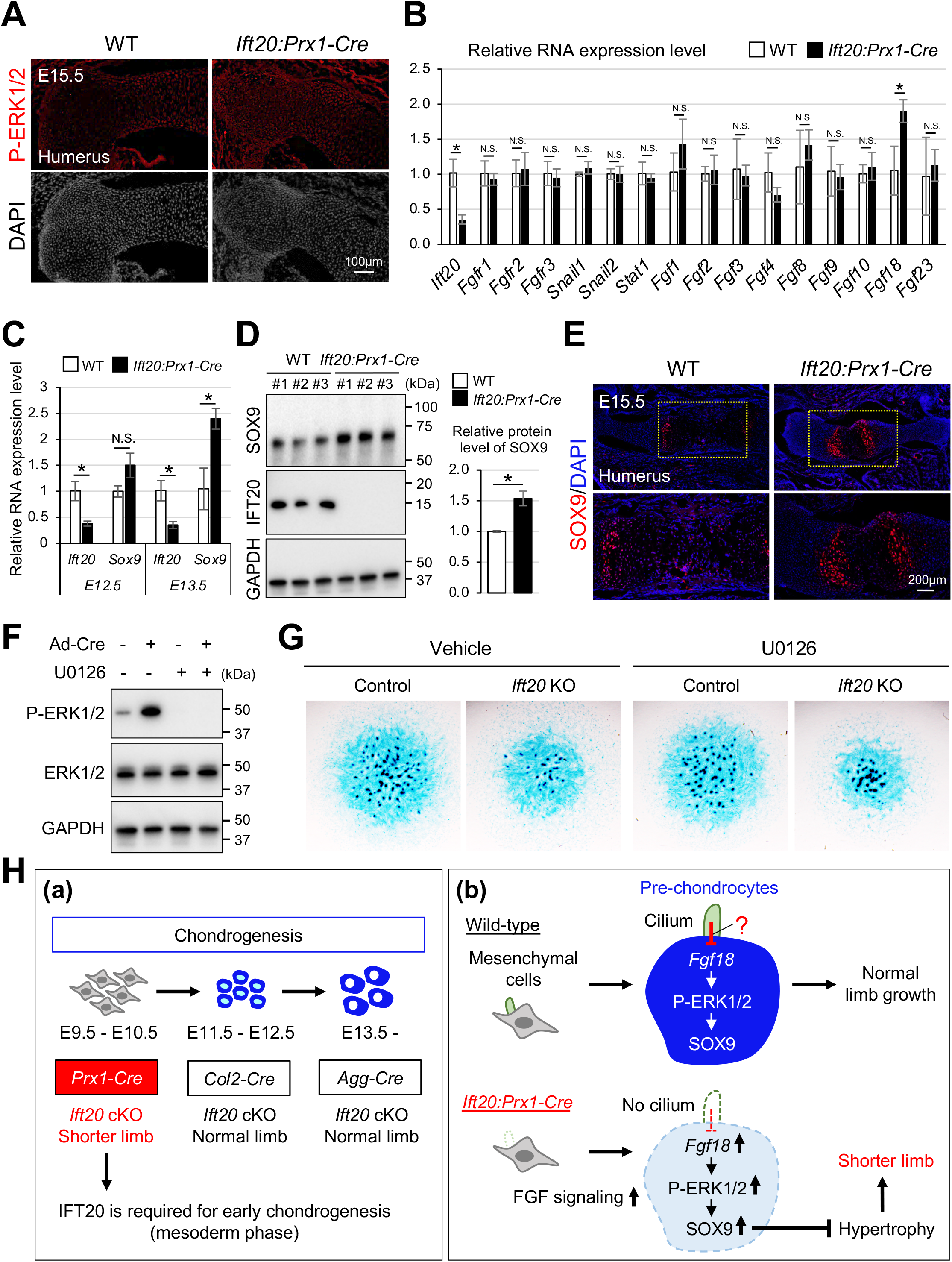
Upregulation of FGF signaling results in abnormal endochondral ossification in *Ift20:Prx1-Cre* mice. **(A)** Immunohistochemical analysis to examine the levels of P-ERK1/2 in humerus. **(B)** qRT-PCR analysis for examining the expression of FGF-related genes using cell lysates from forelimb tissues at E13.5. **(C)** qRT-PCR analysis for examining the expression of *Ift20* and *Sox9* using cell lysates from forelimb tissues. **(D)** Western blot analysis for SOX9 and IFT20 using cell lysates from forelimb tissues at E13.5. Levels of SOX9 production is quantified. **(E)** Immunohistochemical analysis to examine the levels of SOX9. **(F)** Western blotting analysis for p-ERK1/2, ERK1/2 and GAPDH using cell lysates from micromass cultures. **(G)** Alcian blue staining of micromass culture with- or without-U0126 treatment. **(H)** Proposed model for the role of IFT20 in early chondrogenesis during endochondral ossification. **(a)** IFT20 is indispensable for regulating early chondrogenesis. **(b)** In wild-type, primary cilia function restrains FGF signaling pathway. Without cilia, elevation FGF signaling pathway suppresses hypertrophic differentiation via SOX9 sustained expression. This further causes delaying conversion process from chondrocytes and osteoblasts critical for endochondral ossification. Therefore, it leads to shorter limb formation in *Ift20:Prx1-Cre* mice. Data in (B), (C), and (D) are represented as mean ± SD, n = 3 in each group. *p<0.05, N.S., not significant.

At this moment, it is unclear how the deletion of *Ift20* results in the upregulation of *Fgf18*. How does primary cilium restrain *Fgf18* expression? It is noteworthy that another ciliary mutant, Ellis-van Creveld 2 (*Evc2*) mice, displays defective limb outgrowth and shows an elevation of FGF signaling due to the upregulation of *Fgf18*^49^. Therefore, it may be reasonable to speculate that primary cilia may serve as a negative regulator to control FGF signaling during limb outgrowth. This possibility needs to be investigated further in the future.

It has been known that FGF signaling controls the length of cilia^50^. In zebrafish, knockdown of the Fgf receptor and/or Fgf ligand results in shortened primary cilia^51,52^. In mice, a mutation in the FGF receptor causes abnormal ciliogenesis and leads to skeletal defects^53,54^. While FGF signaling is important for regulating cilia length in multiple developmental contexts, our study highlights the importance of ciliary proteins such as IFT20 and Evc2, which are critical for regulating FGF signaling during endochondral ossification.

## Author contributions

Y.K. designed the study. H.Y., M.K., K.U., L.H., J.W., and Y.K. performed the experiments. H.Y., M.K., and Y.K. analyzed the data. D.P., W.L., and Y.K. wrote the manuscript. All authors contributed to the manuscript. Y.K. supervised the project.

## Acknowledgments

We thank Dr. Florent Elefteriou for the *Col2-Cre* mice. We gratefully acknowledge Dr. Mark Corkins for critical reading of this manuscript. This study was supported by a research grant NIDCR/NIH R01DE025897 (Y.K.) and by a fellowship from the Uehara Memorial Foundation (H.Y.).

## Supplemental figure legends

**Supplemental Fig. 1.**
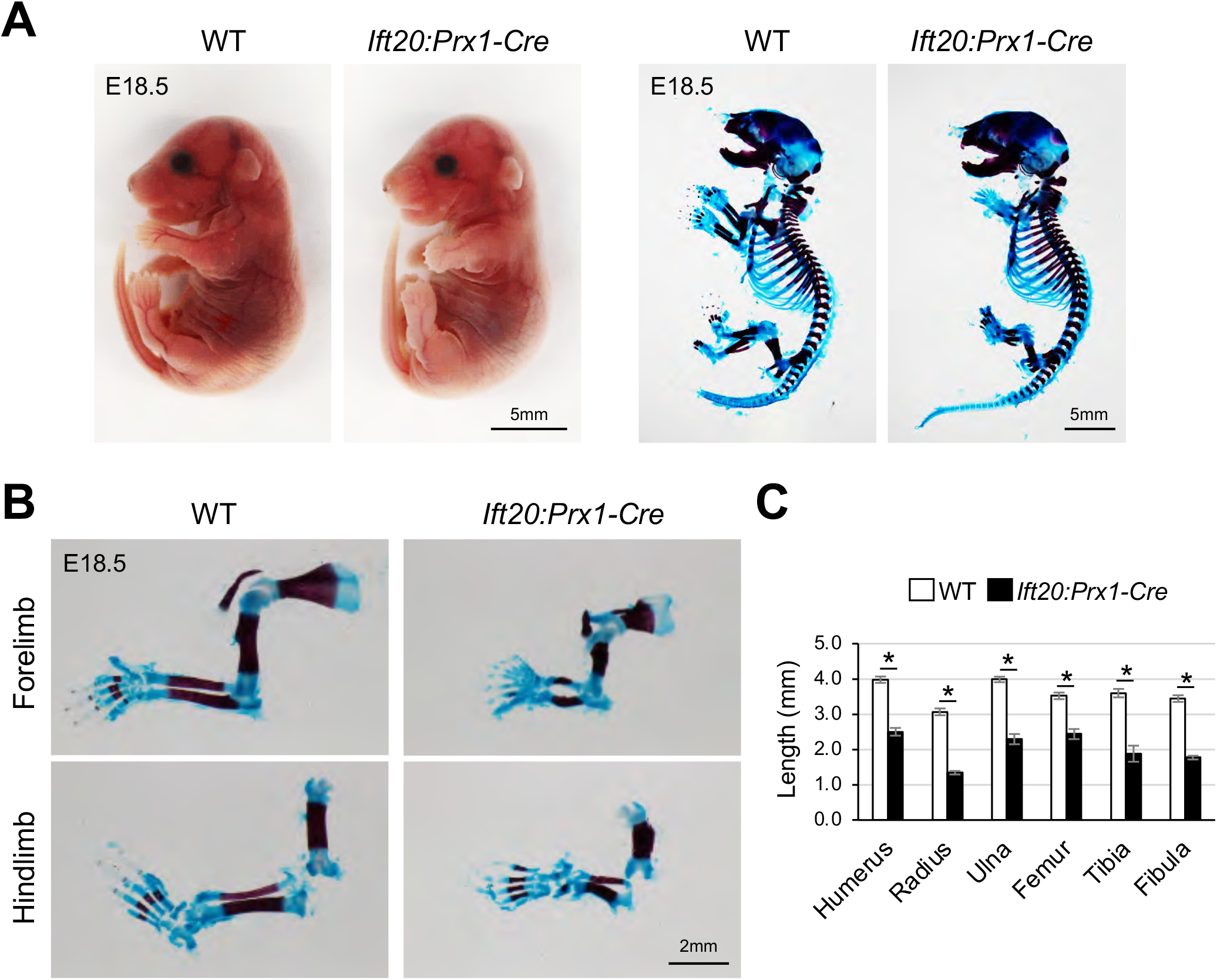
Phenotypic analysis of *Ift20:Prx-Cre* mice. **(A)** Whole mount view and skeletal staining analysis of WT and *Ift20:Prx-Cre* mice. **(B, C)** Skeletal staining and quantification analysis of the length of forelimb and hindlimb in WT and *Ift20:Prx1-Cre* mice at E18.5. Data in (C) are represented as mean ± SD, n = 3 in each group. *p<0.05.

**Supplemental Fig. 2.**
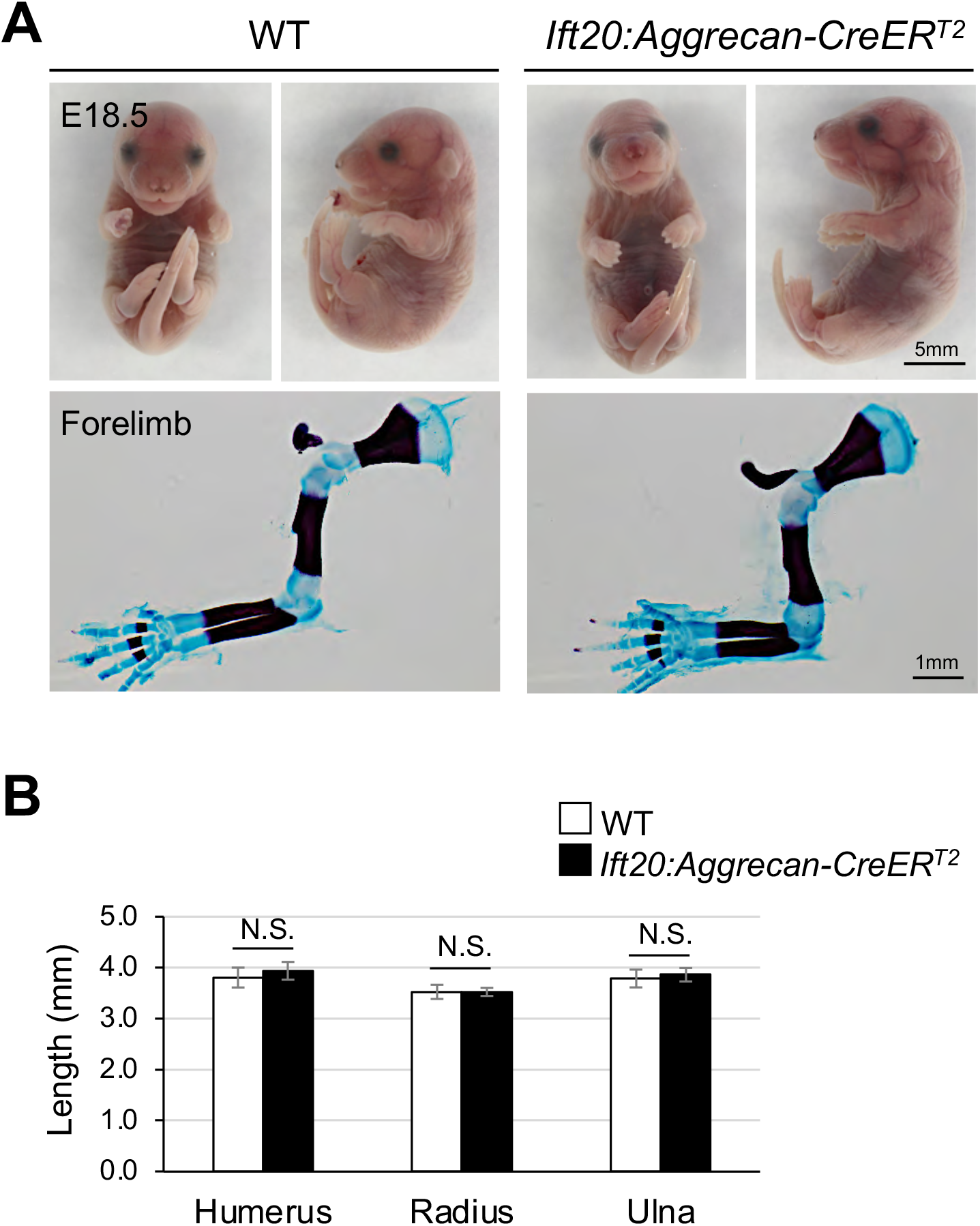
Phenotypic analysis of *Ift20:Aggrecan-CreER^T2^* mice. **(A)** Whole mount and skeletal staining analysis of forelimb from WT and *Ift20:Aggrecan-CreER^T2^* at E18.5. **(B)** Quantification of forelimb length from WT and *Ift20:Aggrecan-CreER^T2^* at E18.5. Data in (B) are represented as mean ± SD, n = 3 in each group. N.S., not significant.

**Supplemental Fig. 3.**
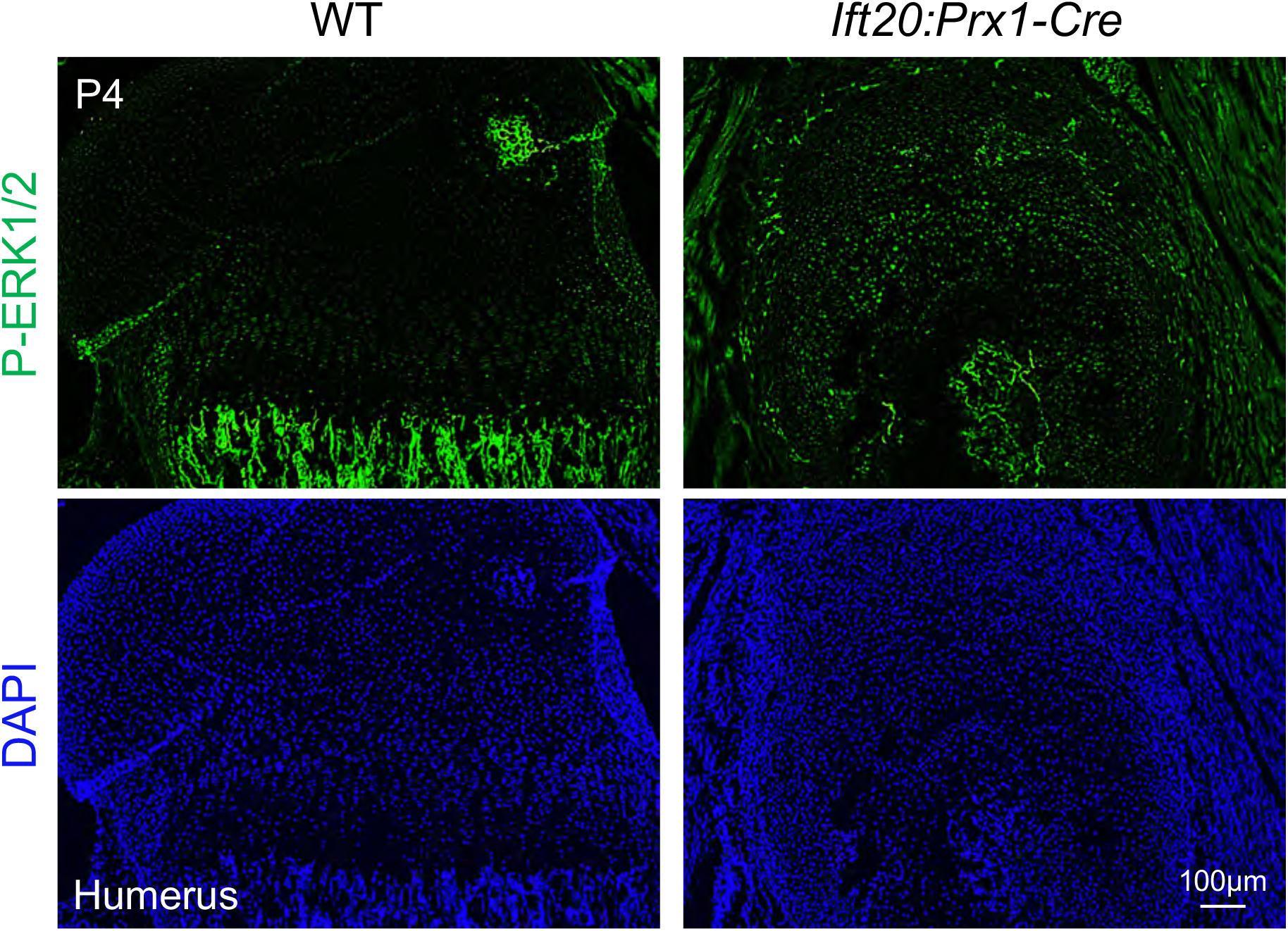
Analysis of p-ERK1/2 in *Ift20:Prx-Cre* mice at postnatal stage. Levels of p-ERK1/2 was examined by immunohistochemistry (green).

**Supplemental Fig. 4.**
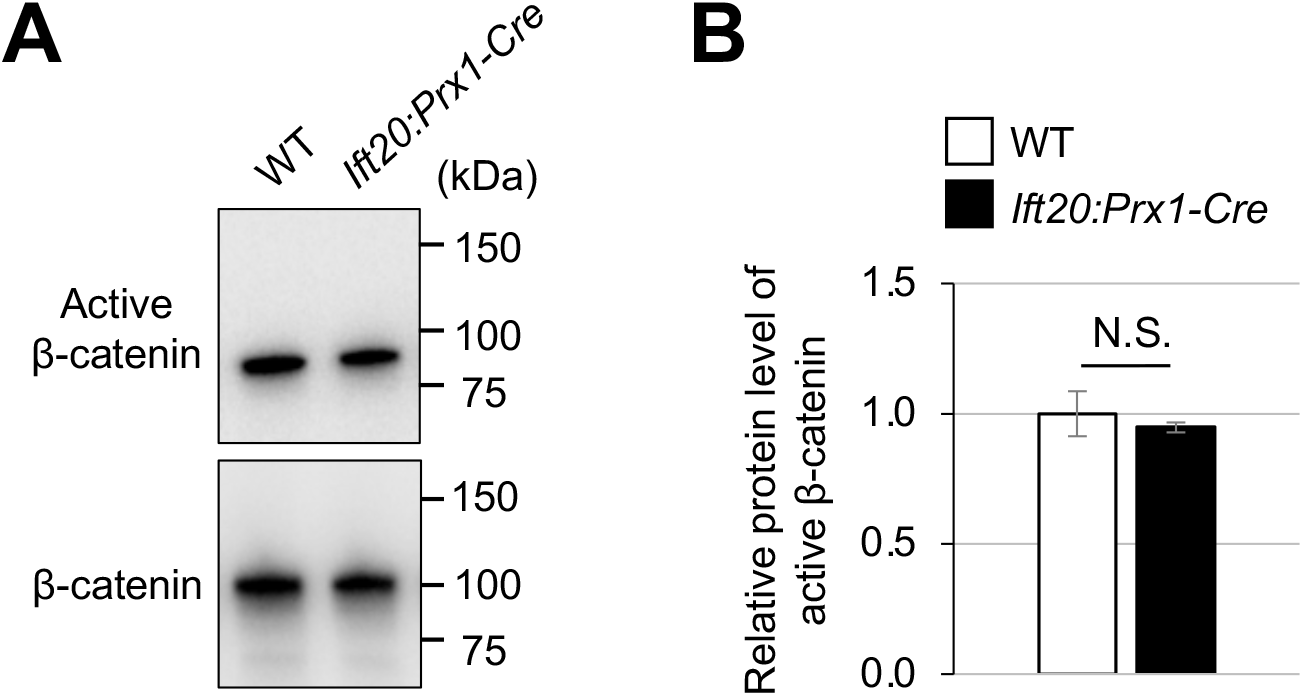
Wnt/β-catenin signaling is intact in *Ift20:Prx-Cre* mice. **(A)** Western blotting analysis to examine active β-catenin and total β-catenin using cell lysates from forelimb tissues at E13.5. **(B)** Quantification analysis of active β-catenin and total β-catenin. Data in (B) are represented as mean ± SD, n = 3 in each group. N.S., not significant.

**Supplemental Table 1.**
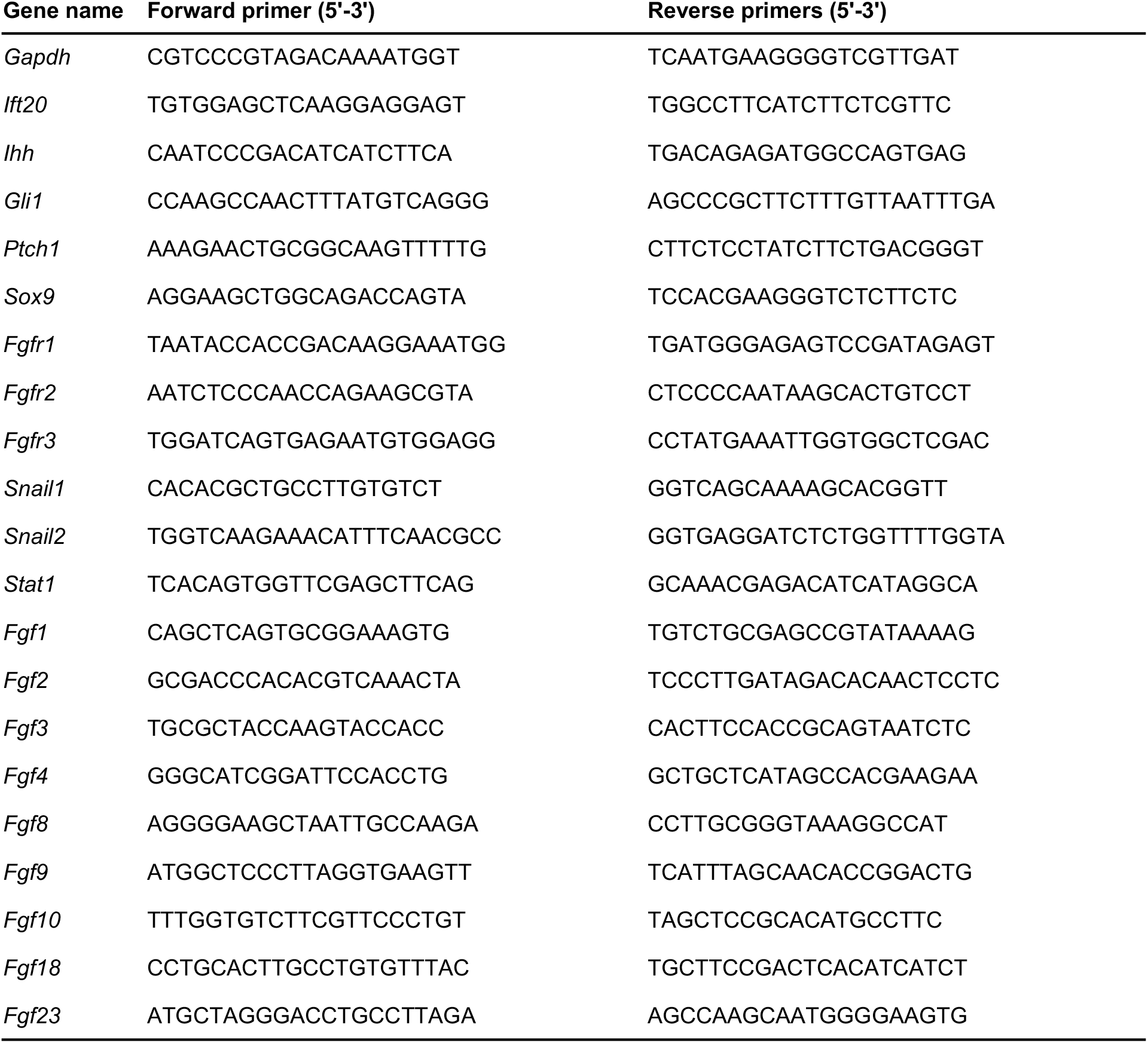
Primer sequences for quantitative RT-PCR used in this study.

**Supplemental video.**
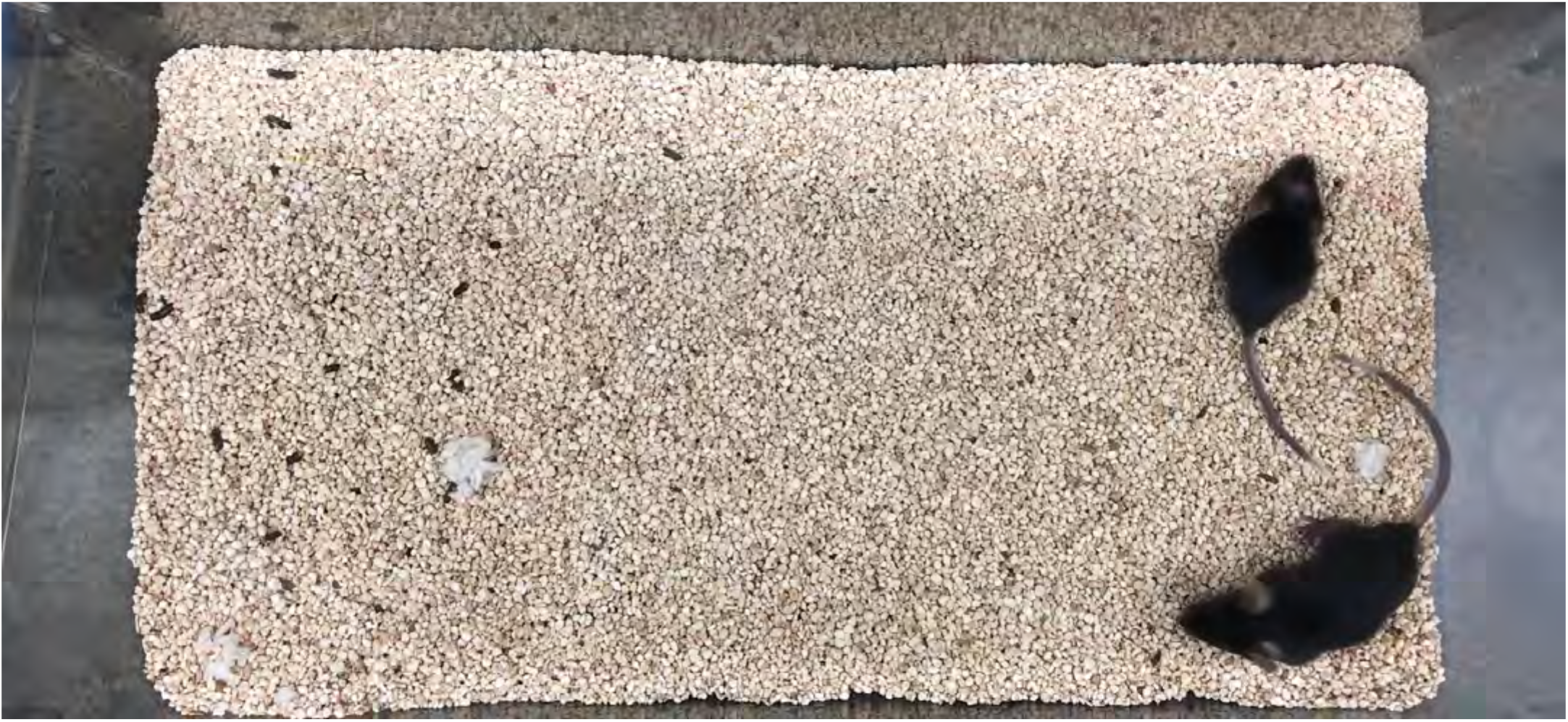
This movie shows the locomotion of WT (large body size) and *Ift20:Prx1-Cre* mice (small body size) at P25.

